# Spatial Transcriptomics Identifies Muscle Inflammation Susceptibility as a Distinct Periarticular Skeletal Muscle Phenotype in End-Stage Knee Osteoarthritis

**DOI:** 10.64898/2026.09.16.752138

**Authors:** Franccesco P. Boeno, Zachary A. Graham, Qi Su, Michael J. Stec, Arny A. Ferrando, Wei Keat Lim, Marcas M. Bamman

## Abstract

Skeletal muscle dysfunction is a major contributor to disability and incomplete functional recovery in patients with end-stage knee osteoarthritis (KOA), yet the spatial components of disease-associated molecular remodeling remain poorly understood. Here, we applied spatial transcriptomics to paired skeletal muscle biopsies obtained from the surgical (Sx) and contralateral (Ct) limbs of individuals undergoing total knee arthroplasty (TKA) to define the cellular architecture of periarticular muscle and determine how muscle inflammation susceptibility (MuIS) shapes local transcriptional programs. Integrated analysis of 27,087 spots obtained from 22 muscle histological cross-sections (11 Sx-Ct pairs) revealed seven spatially resolved transcriptional domains corresponding to slow and fast myofiber states, an extracellular matrix/fibroadipogenic-enriched domain, a pericyte/smooth muscle domain, and a satellite cell/myogenic-enriched domain. Despite advanced unilateral disease, the major annotated cellular compartments were similarly represented between Sx and Ct limbs. KOA-associated remodeling was reflected primarily by within-cluster transcriptional changes, with the most informative differences observed in fibroadipogenic, pericyte/smooth muscle, and satellite/myogenic domains. Within Sx, MuIS stratification identified a coordinated transcriptional program characterized by denervation-and regeneration-associated genes and altered contractile and metabolic features. Neighborhood analysis localized denervation-associated signals primarily to fast-myofiber-rich regions, while local adjacency patterns among the examined myofiber, fibroadipogenic, and pericyte/smooth muscle domains were broadly preserved. Our findings provide the first spatial transcriptomic analysis of periarticular skeletal muscle in end-stage KOA and identify MuIS as a distinct local transcriptional phenotype in diseased muscle.

## INTRODUCTION

Osteoarthritis (OA) is the most common form of arthritis worldwide and a leading cause of pain, disability, and reduced physical function^1,2^. OA is traditionally viewed as a disease of the joint, yet functional recovery after total knee arthroplasty (TKA) is strongly influenced by the health of the periarticular skeletal muscle^3,4^. Patients with knee OA (KOA) commonly exhibit quadriceps weakness, impaired mobility, and reduced muscle quality, and these deficits may persist after TKA^5^. Thus, skeletal muscle is increasingly recognized as an active determinant of disability and recovery in end-stage KOA rather than a passive consequence of joint degeneration^6,7^. Via whole tissue transcriptomic profiling, we recently identified substantial differences in the gene expression profile of muscle from periarticular muscle samples from surgical (Sx) vs. contralateral (Ct) thighs, indicative of inflammation, pro-fibrosis, lipid metabolism, and overall poor muscle quality on the Sx limb^7^.

Over the past decade, our group has progressively characterized a distinct biological phenotype in a subset of individuals termed muscle inflammation susceptibility (MuIS)^8–10^. Initially described in aging skeletal muscle, MuIS was defined as an intrinsic propensity for exaggerated inflammatory signaling that is localized to skeletal muscle and not detected systemically, resulting in impaired myogenesis and regenerative capacity^8^. This concept was subsequently translated to patients undergoing total hip arthroplasty, where MuIS+ individuals expressed a localized periarticular inflammatory muscle phenotype independent of systemic inflammation, associated with impaired anabolic signaling and muscle remodeling^9^. More recently, we refined MuIS in patients with end-stage KOA by demonstrating that elevated Sx muscle TNF-α receptor (TNF-αR) expression identifies MuIS+ individuals who show heightened inflammatory and catabolic gene expression and a trend toward greater fibrosis^10^.

Despite these advances, the cellular resolution of periarticular skeletal muscle in KOA, and the transcriptional programs underlying MuIS, remain unknown. Previous studies relied primarily on bulk gene-expression profiling of periarticular muscle in OA and MuIS^7,11,12^, which cannot distinguish whether transcriptional circuits arise from myofibers, fibroadipogenic progenitors (FAPs), pericytes, endothelial cells, satellite cells, immune populations, or coordinated interactions among these resident niches. Because skeletal muscle is a highly organized but heterogeneous tissue in which contractile, stromal, vascular, and regenerative compartments communicate extensively to regulate tissue homeostasis^3,13^, resolving the anatomical context of periarticular muscle in KOA is essential for understanding the multicellular basis of MuIS and the biological mechanisms underlying muscle dysfunction.

Spatial transcriptomics addresses this limitation by preserving tissue architecture while resolving gene-expression programs within intact sections^14^. In skeletal muscle, this approach can localize transcriptional states to discrete cellular compartments that are obscured by conventional bulk RNA sequencing. Recent applications of spatial transcriptomics from our group^15^ and others^16–18^ have demonstrated regional organization and coordinated cellular responses in human skeletal muscle^15,18^, while applications in inflammatory joint tissues have revealed spatially restricted immune–stromal interactions that cannot be inferred from tissue homogenates^17^. However, spatial transcriptomics has not previously been applied to human periarticular skeletal muscle in KOA.

Here, we performed spatial transcriptomic profiling of paired skeletal muscle biopsies collected from the surgical and contralateral limbs of individuals with end-stage KOA undergoing TKA. We first characterized the spatial cellular architecture of periarticular skeletal muscle and determined how it differs between limbs. We then investigated how MuIS influences transcriptional programs within individual spatial domains while preserving tissue organization. We hypothesized that MuIS would be characterized by coordinated transcriptional remodeling across myofiber and interstitial compartments of skeletal muscle.

## METHODS

### Experimental model and subject details

#### Human participants

This analysis focused on a subset of 11 individuals with end-stage KOA undergoing TKA who were recruited at the University of Alabama at Birmingham (UAB) and the University of Arkansas for Medical Sciences (UAMS) as part of a parent clinical trial (NCT02628795). In the parent trial, participants were excluded for conditions or treatments that could influence skeletal muscle biology beyond end-stage OA or preclude participation in the study, including bilateral arthroplasty, uncontrolled cardiometabolic disease, recent resistance training, use of anabolic agents, or other investigator-determined exclusion criteria. The full description of the study criteria was previously described^7,10^. All participants provided written informed consent for future use of biospecimens. The study was approved by the Institutional Review Boards of UAB and UAMS and conducted in accordance with the Declaration of Helsinki.

#### Muscle samples

Paired skeletal muscle biopsies were obtained from the surgical limb (Sx) and the contralateral nonsurgical limb (Ct) from each participant, enabling within-subject comparison of local disease burden and between-group comparison based on MuIS status. Participants underwent biopsy collection in the fasting state according to the TKA scheduling. For the contralateral limb, muscle biopsy samples were obtained from the vastus lateralis using a 5-mm Bergström needle with suction immediately before surgical incision as previously described^19^ after surgical-plane anesthesia was established. For the surgical limb, muscle samples were obtained from the vastus medialis using a scalpel during the arthroplasty procedure. Samples were immediately trimmed to remove visible connective tissue, adipose tissue, and blood^10^. A portion of each biopsy (∼50 mg) was placed in a mixture of Optimal Cutting Temperature compound and tragacanth gum and subsequently frozen in liquid nitrogen cooled isopentane for histological analyses. Histological mounts were shipped to Regeneron overnight on dry ice for sectioning and downstream spatial gene expression profiling using the standard 10x Genomics Visium CytAssist workflow.

#### Muscle inflammation susceptibility (MuIS)

MuIS status was previously determined for each participant based on TNF-αR gene expression in the vastus medialis of the Sx limb as described^10^. Participants were classified as either muscle inflammation susceptible (MuIS+) or non-susceptible (MuIS-) according to our previously established perioperative profiling strategy using unbiased K-means clustering^10^ similar to our approach in other studies^20^. The cluster with the lowest TNF-αR receptor expression was designated MuIS-whereas the remaining participants with higher expression were designated MuIS+.

#### Thigh Muscle Mass and WOMAC assessment

Preoperative thigh lean mass of Sx and Ct limbs was assessed by dual-energy X-ray absorptiometry (DXA) following a previously validated protocol^10^ prior to the surgery. Thigh lean mass was normalized to participant height (m) and transformed to kg/m^2^. Preoperative OA-related pain, stiffness, and physical function were assessed using the Western Ontario and McMaster Universities Osteoarthritis Index (WOMAC), a validated patient-reported outcome measure completed before surgery^21^.

### Spatial transcriptomics analysis

#### Spatial transcriptomics tissue processing and library preparation

Frozen skeletal muscle samples were cryosectioned and mounted according to the Visium CytAssist spatial gene expression workflow for fresh frozen tissue. Tissue sections were immunostained for fiber typing as previously described^15^. Briefly, samples were fixed with ice-cold methanol, blocked, incubated with primary antibodies (Type I, DSHB Cat #BA-F8; Type IIa, DSHB Cat #SC-71; Laminin rabbit; Sigma Aldrich Cat #L9393), washed, incubated with secondary antibodies (mIgG2b 546, Invitrogen Cat #A21143; mIgG1 647, Invitrogen Cat #A21240; rIgG 488, Invitrogen Cat #A32731; Hoechst 33342, Invitrogen Cat #H3570), washed, immersed in 3x SSC buffer, and mounted with glycerol. Slides were immediately imaged using a Zeiss Axioscan Z1, and then further processed for spatial transcriptomics. Cross section area (CSA) and fiber typing analysis were performed using Myovision 2.0 software. Following imaging, transcriptomic probes were hybridized to tissue sections and transferred to Visium slides containing spatially barcoded capture spots with a nominal spot diameter of 55 μm. Library preparation and sequencing were performed according to the manufacturer’s protocol (10x Genomics Visium CytAssist). Raw sequencing data (FASTQ files) generated for each sample were processed using the Space Ranger to perform sample demultiplexing, read alignment, tissue detection, and generation of gene expression counts.

#### Spatial transcriptomics data loading and quality control

Spatial count matrices were imported into Seurat using Load10X_Spatial. Sample names were parsed to capture participant ID, limb group (Ct or Sx), and inflammatory classification (MuIS+ or MuIS-). Quality control was performed at the spot level. For each sample, the number of detected genes per spot (nFeature_Spatial), the number of UMI counts per spot (nCount_Spatial), and the percentage of mitochondrial transcripts were calculated. Spots were retained if they met the following thresholds: more than 200 detected genes per spot and more than 500 UMI counts per spot. No explicit upper threshold was applied for gene or UMI counts, and no mitochondrial-content cutoff was used. Quality control summaries and per-sample filtering metrics were generated prior to downstream analysis.

#### Normalization, integration, dimensionality reduction, and clustering

Each sample was first normalized using SCTransform in Seurat. Shared variable features were identified across samples, and the most recurrent features were used to define the integration feature set. SCT integration was then performed per sample using PrepSCTIntegration, FindIntegrationAnchors, and IntegrateData in Seurat. The integrated dataset was analyzed using principal-component analysis (PCA) followed by uniform manifold approximation and projection (UMAP). Neighbor finding and unsupervised graph-based clustering were performed using the first 10 principal components with clustering resolution set to 0.4. The integrated object was then used for visualization, cluster annotation, cluster composition analysis, and differential expression testing.

#### Cluster annotation

Clusters were annotated based on canonical marker gene expression using a combination of marker discovery, average-expression profiles, and prior biological knowledge. Cluster markers were identified using FindAllMarkers on the SCT assay after PrepSCTFindMarkers, with only.pos = TRUE, min.pct = 0.1, and logfc.threshold = 0.1. Canonical marker genes representing major skeletal muscle and stromal cell states were visualized using dot plots and heatmaps.

#### Cluster composition analysis

Cluster composition was quantified at the sample level as the proportion of spots assigned to each annotated cluster relative to the total number of retained spots in that sample. For the paired Ct vs. Sx comparison, proportions were compared using paired Wilcoxon signed-rank tests. For MuIS+ vs. MuIS-comparisons within the Sx- and Ct-only integration, group differences in cluster proportions were assessed using Wilcoxon rank-sum tests.

#### Differential expression analysis

Within-cluster differential expression was performed using Seurat on the SCT assay after PrepSCTFindMarkers. For the integrated Ct vs. Sx analysis, spots were compared within each annotated cluster between Sx and Ct samples. For the Sx- and Ct-only analysis, spots were compared within each cluster between MuIS+ and MuIS- groups. Differential expression was calculated using Seurat FindMarkers with default Wilcoxon rank-sum testing unless otherwise specified. Genes with adjusted p value < 0.05 were considered statistically significant for downstream interpretation and enrichment analyses. We further applied a threshold of adjusted p value < 0.05 and absolute log2 fold-change ≥ log2(1.5) to emphasize more robust expression differences.

#### Functional enrichment analysis

Functional enrichment was performed using clusterProfiler with human gene annotation from org.Hs.eg.db. Direction-specific gene sets were analyzed separately, and KEGG results were visualized with dot plots in which dot size represented gene count and color represented enrichment significance as −log10 (adjusted p value).

#### Sx- and Ct-only MuIS reintegration and focused transcriptional analysis

To examine transcriptional differences associated with inflammatory susceptibility in the diseased limb, Sx and Ct samples were reintegrated and analyzed independently. The same SCTransform-based normalization, integration, dimensionality reduction, and clustering framework described above was applied to both subsets. MuIS status was carried forward from the TNF-αR-based stratification. Differential expression and pathway enrichment were then performed within each Sx-and Ct-derived cluster comparing MuIS+ and MuIS-samples.

#### Spatial neighborhood and hotspot analysis

Neighborhood analysis was performed on the Sx-only integrated dataset using the annotated fast myofiber 1, fibroadipogenic, and pericyte/smooth muscle clusters. Cluster identities were projected back onto tissue coordinates to generate reconstructed spot-based maps and cluster outlines. Neighborhood enrichment was quantified by comparing the observed frequency of adjacency between cluster pairs with the frequency expected under random spatial arrangement, generating neighborhood z-scores for each cluster-to-cluster interaction within each MuIS group. Positive z-scores indicate enriched spatial adjacency, whereas negative z-scores indicate relative spatial separation. For hotspot analysis, a composite denervation-associated score was generated from the spatial expression of MYH8, CHRNG, CHRNA1, and MUSK. These genes were selected from the MuIS+ versus MuIS-differential expression analysis in the fast myofiber 1 cluster based on their strong enrichment and known association with denervation and regenerative myofiber remodeling. Local hotspot scores were calculated from the combined expression of these genes and overlaid onto reconstructed tissue maps for spatial visualization.

#### Clinical phenotype integration

Clinical phenotype integration was performed using participant-level body composition and symptom data. DXA-derived normalized lean mass variables from the surgical and contralateral leg and thigh were aligned to the subset of participants included in the spatial transcriptomic dataset. WOMAC total and physical function scores were also harmonized to the same participant IDs. For each subject, cluster proportion was calculated as the fraction of Sx spots assigned to a given cluster. In addition, subject-level pseudobulk expression matrices were generated within each cluster by summing expression across all spots assigned to that cluster for each subject. MuIS-associated cluster signature scores were then derived as the mean z-scored expression of the top genes distinguishing MuIS+ and MuIS- within that cluster. Associations between cluster-level spatial features and clinical phenotypes were evaluated using Spearman rank correlation.

#### Quantification and statistical analysis

All analyses were performed in R using Seurat, ggplot2, dplyr, patchwork, clusterProfiler, and related packages. Unless otherwise stated, statistical significance was assessed at adjusted p < 0.05 for gene-level and pathway-level analyses. For paired Ct vs. Sx cluster composition analyses, Wilcoxon signed-rank tests were used. For unpaired MuIS+ vs. MuIS-comparisons, Wilcoxon rank-sum tests were used. Correlation analyses were performed using Spearman’s rho. Multiple-testing adjustment for differential expression and enrichment analysis was performed using the procedures implemented in Seurat and clusterProfiler.

### Data and code availability

Spatial transcriptomic data generated, processed count matrices, cluster annotations, differential expression results, and summary tables that are not available in the supplemental materials can be requested from the lead contact author upon reasonable request.

## RESULTS

Spatial transcriptomic profiling was performed on paired Ct and Sx skeletal muscle biopsies obtained from 11 participants undergoing TKA, including 5 classified as MuIS+ and 6 as MuIS-(**Fig 1A**). The two groups were similar in age, normalized leg and thigh lean mass, and WOMAC-derived symptom measures, with no significant baseline differences detected between groups (Table 1). Mean age was 62.6 ± 3.8 years in the MuIS+ group and 61.4 ± 11.9 years in the MuIS-group. Sex distribution was uneven across groups, with only one woman represented in the MuIS+ group, although the cohort was not powered for sex-stratified analysis. **Table 1** shows demographics and sample characterization. We also performed myofiber morphometric analysis of the paired biopsies showed no significant differences in fiber area or fiber type proportion between Ct and Sx muscles. In contrast, fiber shape analysis identified higher median shape index values in the surgical limb for all fibers (p = 0.049) and for type 2 fibers (p = 0.035), indicating a less regular and more angular fiber profile in Sx muscle (Table 2). No differences were found between MuIS+ and MuIS-in either the Ct or Sx limb.

**Figure 1.**
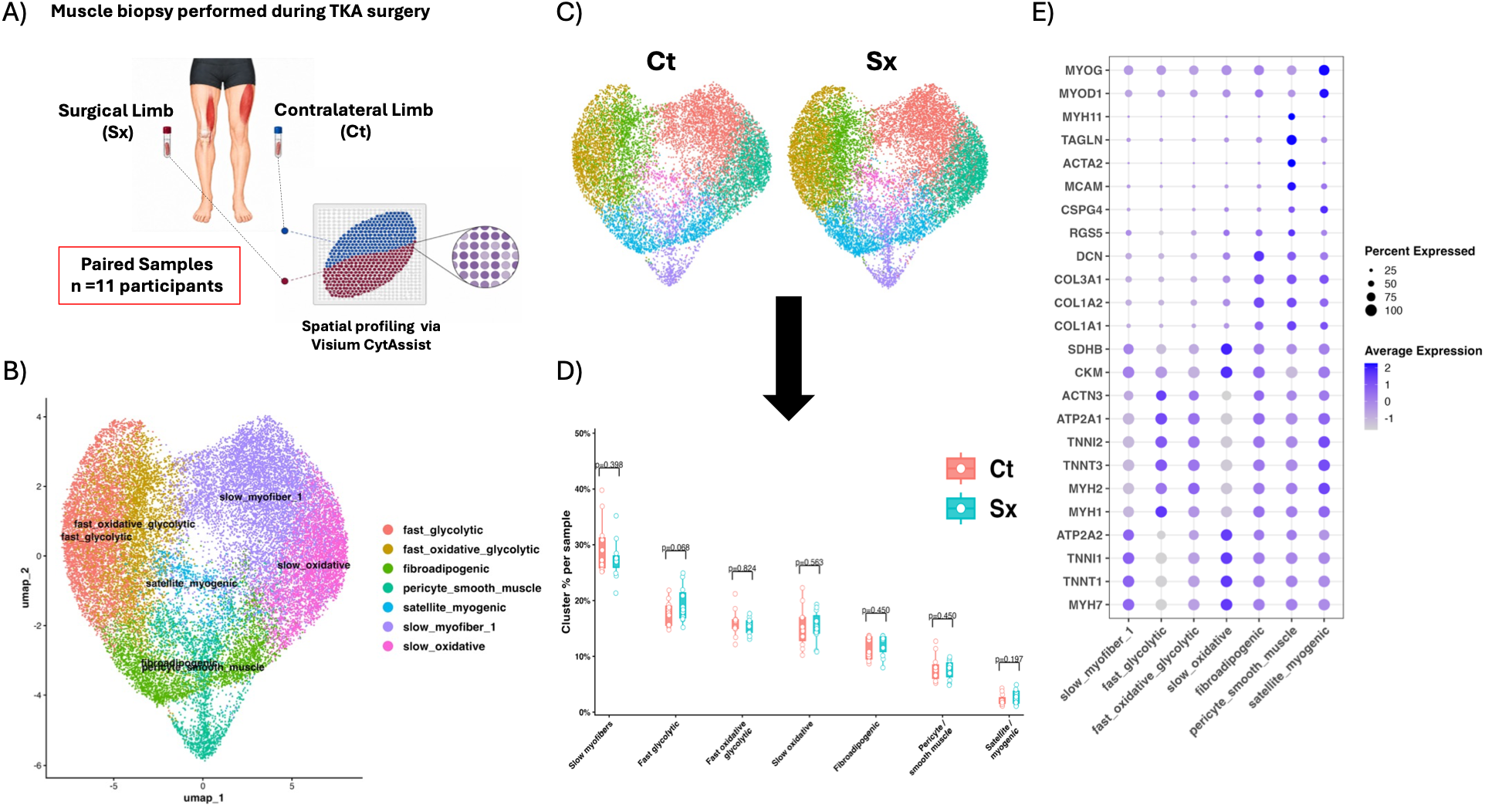
Integrated spatial transcriptomic analysis identifies major skeletal muscle cell states across TKA samples. A) study design. B) UMAP of all quality-controlled spatial spots following integration of all samples, colored by annotated cluster identity. Unsupervised clustering resolved seven major transcriptional domains corresponding to fast glycolytic myofibers, fast oxidative-glycolytic myofibers, slow myofibers, slow oxidative myofibers, fibroadipogenic cells, pericyte/smooth muscle cells, and satellite/myogenic cells. C) Split UMAPs showing the distribution of annotated clusters in contralateral control (Ct) and surgical (Sx) muscle samples. D) Cluster composition per sample in Ct and Sx muscle, shown as the percentage of spots assigned to each cluster. Points represent individual samples, and p values indicate Wilcoxon signed-rank tests comparisons between Ct and Sx samples. E) Dot plot of canonical marker genes used for cluster annotation.

**Table 1.** Participant characteristics. Baseline demographic, body composition, and clinical characteristics of participants included in the spatial transcriptomic analysis, grouped by MuIS+ and MuIS-status. Values are presented as mean ± SD unless otherwise indicated. DXA-derived lean mass variables were normalized by patient’s height (m). Clinical status was assessed using the Western Ontario and McMaster Universities Arthritis Index (WOMAC) with pre-surgical total and physical function scores highlighted.

|  | MuIS+ (N=5) | MuIS- (N=6) | All participants (11) |
| --- | --- | --- | --- |
| Age (y) | 62.6 ± 3.8 | 61.4 ± 11.9 | 62 ± 8.4 |
| Sex (M/F) | 4/1 | 3/3 | 7/4 |
| Contralateral leg lean mass (kg/m<sup>2</sup>) | 2.7 ± 0.19 | 2.8 ± 0.20 | 2.8 ± 0.2 |
| Surgical leg lean mass (kg/m<sup>2</sup>) | 2.6 ± 0.33 | 2.8 ± 0.22 | 2.2 ± 0.3 |
| Contralateral thigh lean mass (kg/m<sup>2</sup>) | 1.9 ± 0.12 | 1.9 ± 0.19 | 1.9 ± 0.2 |
| Surgical thigh lean mass (kg/m<sup>2</sup>) | 1.82 ± 0.18 | 1.97 ± 0.24 | 1.91 ± 0.2 |
| WOMAC total score | 59 ± 21 | 45 ± 25 | 54 ± 21 |
| WOMAC physical function score | 38.2 ± 12.9 | 29 ± 18.4 | 35.6 ± 13.7 |

**Table 2.** Myofiber morphometric and shape index characteristics in contralateral and surgical muscle. Values are presented as median [IQR]. P values were calculated using paired t tests comparing contralateral (Ct) and surgical (Sx) muscles within subject. Ct muscle samples are derived from *vastus lateralis* and Sx samples from v*astus medialis*.

|  | Contralateral Limb<br>(VL) | Surgical Limb<br>(VM) | Paired t-test<br>p-value |
| --- | --- | --- | --- |
| All fibers, Median Area (um<sup>2</sup>) | 2876 [2331, 3523] | 4069 [2487, 4274] | 0.145 |
| Type 1 Fibers, Median Area (um<sup>2</sup>) | 2981 [2625, 3893] | 4318 [3054, 4855] | 0.161 |
| Type 2 Fibers, Median Area (um<sup>2</sup>) | 2246 [1984, 3516] | 3933 [2237, 4198] | 0.090 |
| Type 1 Fiber Proportion | 0.46 [0.30, 0.54] | 0.35 [0.32, 0.46] | 0.383 |
| Type 2 Fiber proportion | 0.54 [0.46, 0.70] | 0.65 [0.54, 0.68] | 0.383 |
| All Fibers, Median Shape Index | 1.44 [1.42, 1.50] | 1.52 [1.48, 1.55] | 0.049 |
| Type 1 Fibers, Median Shape Index | 1.38 [1.35, 1.48] | 1.44 [1.40, 1.48] | 0.407 |
| Type 2 Fibers, Median Shape Index | 1.50 [1.45, 1.53] | 1.58 [1.53, 1.62] | 0.035 |

### Sx and Ct Muscles Share Major Spatial Domains but Differ in Within-Cluster Transcriptional Programs

The average number of probe spots analyzed per sample was 1,231 ± 474, and the median reads per spot was 18,196 (IQR 9,705-27,298) (**Table S1**). Following quality control, we performed integration of Sx and Ct samples to generate a unified cellular atlas and enable direct comparison of shared cell populations across Sx and Ct muscles. Unsupervised clustering identified multiple transcriptionally distinct spatial domains across skeletal muscle sections. UMAP analysis resolved 27,087 spatial spots into 7 major clusters. Annotation using canonical marker genes identified clusters corresponding to (1) slow myofibers, (2) fast glycolytic myofibers, (3) fast oxidative-glycolytic myofibers, (4) slow oxidative myofibers, (5) fibroadipogenic stromal cells, (6) pericyte/smooth muscle cells, and (7) satellite/myogenic cells (**Fig 1B**).

Slow myofiber clusters were defined by enrichment of MYH7, TNNT1, and TNNI1, whereas fast myofiber clusters were defined by enrichment of MYH1, MYH2, TNNT3, TNNI2, and ATP2A1. The slow myofiber cluster expressed a canonical slow-fiber program, whereas the slow oxidative cluster showed a more oxidative slow-fiber state, characterized in part by HMGCS2, OXCT1, and LDHB. The fibroadipogenic cluster was characterized by extracellular matrix-associated genes including COL1A1, COL1A2, COL3A1, DCN, LUM, and PDGFRA, while the pericyte/smooth muscle cluster expressed RGS5, CSPG4, MCAM, ACTA2, TAGLN, and MYH11. A smaller satellite/myogenic cluster was identified by expression of PAX7, MYOD1, MYOG, and NCAM1 (**Fig 1E**). Among the integrated clusters, slow myofiber and fast myofiber populations comprised the majority of spatial spots. The largest cluster contained 7,625 spots and corresponded to a slow myofiber state, followed by clusters containing 5,065 and 4,254 spots representing fast glycolytic and fast oxidative-glycolytic myofibers, respectively. Additional clusters included 4,232 spots assigned to slow oxidative myofibers, 3,197 spots assigned to fibroadipogenic cells, 2,050 spots assigned to pericyte/smooth muscle cells, and 664 spots assigned to satellite/myogenic cells. Cluster proportions did not differ significantly between limbs for any of the seven domains (all p ≥ 0.068 by paired Wilcoxon signed-rank; **Fig 1D**), with the smallest p-value observed for the fast glycolytic cluster (p = 0.068). Within-cluster differential expression analysis comparing Sx vs. Ct identified transcriptional remodeling across all clusters. The largest number of differentially expressed genes (DEG) was observed in satellite/myogenic cells (n = 137), followed by slow myofiber 1 (n = 130), fast glycolytic (n = 126), fast oxidative_glycolytic (n = 126), fibroadipogenic (n = 107), slow oxidative (n = 107), and pericyte/smooth muscle cells (n = 95). The clusters highlighted in **Fig 2** were prioritized because they capture biologically relevant non-myofiber niches involved in stromal remodeling, vascular support, and regenerative signaling.

**Figure 2.**
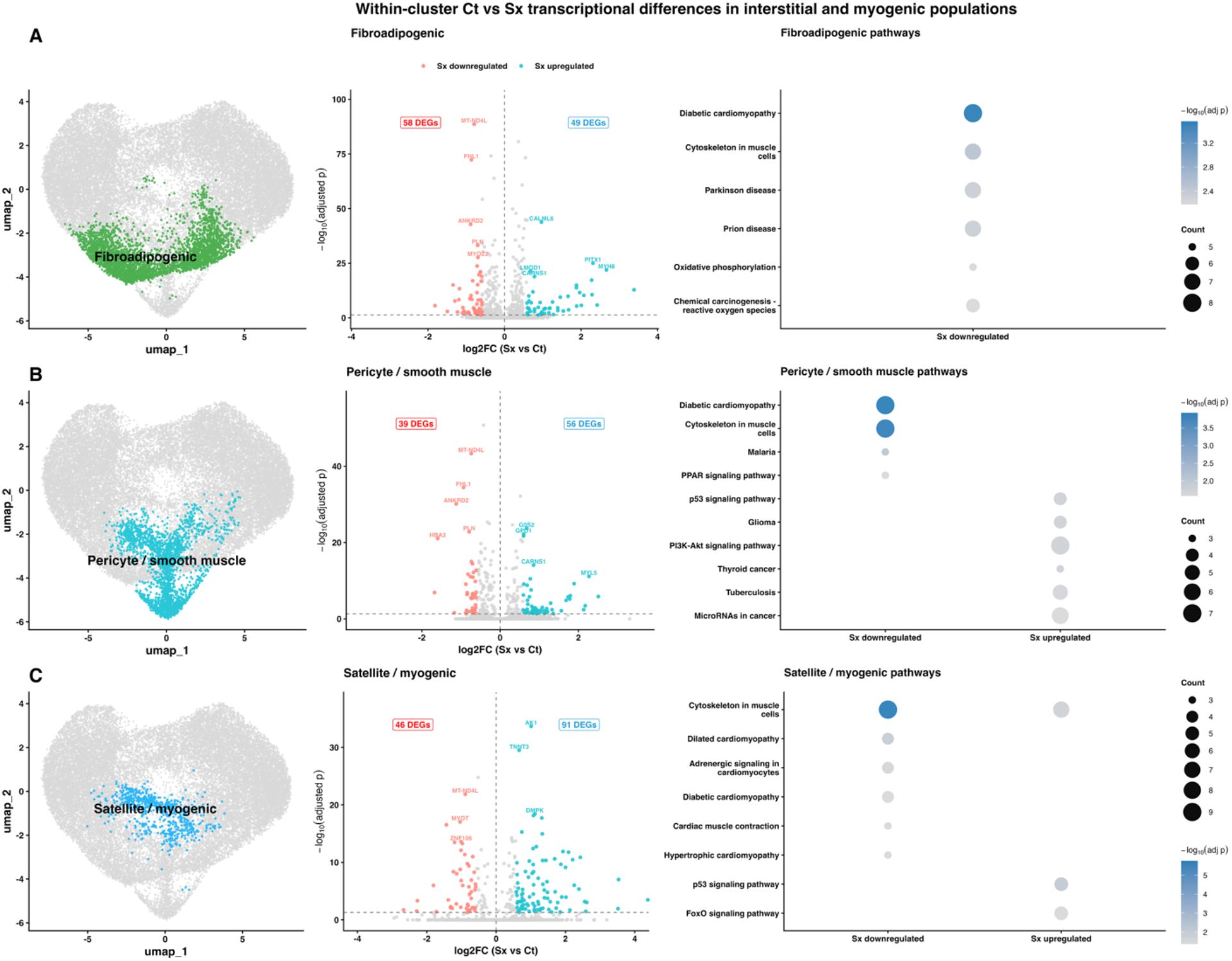
Within-cluster transcriptional differences between contralateral and surgical muscle in interstitial and myogenic populations. Differential gene expression and pathway enrichment analyses comparing surgical (Sx) and contralateral control (Ct) muscle within A) fibroadipogenic, B) pericyte/smooth muscle, and C) satellite/myogenic clusters from the fully integrated dataset. Left panels show volcano plots of spot-level gene expression changes for Sx relative to Ct, with positive log2 fold-change indicating genes upregulated in Sx and negative log2 fold-change indicating genes downregulated in Sx. Selected top genes meeting significance criteria are annotated. Right panels show KEGG pathway enrichment results derived from high-confidence differentially expressed genes, defined as adjusted p value < 0.05 and absolute log2 fold-change of at least log2 (1.5). Dot size indicates the number of genes contributing to each pathway, and color intensity reflects enrichment significance as −log10 (adjusted p value).

In satellite/myogenic cells, the top genes upregulated in Sx included LTK, ERBB3, PVALB, ANXA1, and DCLK1, known calcium-handling and stress-responsive genes. The top genes downregulated in Sx included TECRL, HBA2, RSPO3, ABRA, and SEMA3C, indicating reduced expression of genes linked to metabolic, cytoskeletal, and regenerative support functions (e.g., RSPO3-linked WNT/β-catenin signaling that supports myogenic differentiation). KEGG analysis further show that genes downregulated in Sx were enriched for cytoskeleton in muscle cells, cardiac muscle contraction, dilated cardiomyopathy, and motor proteins, whereas genes upregulated in Sx were enriched for cytoskeleton in muscle cells, glucagon signaling, and p53 signaling.

In the fibroadipogenic cluster, the top genes upregulated in Sx included ERBB3, MYH8, PVALB, PITX1, and SCT, whereas the top genes downregulated in Sx included HBB, AKR1C3, HBA2, TECRL, and CYP2J2. In pericyte/smooth muscle cluster, the top genes upregulated in Sx included ERBB3, MYL5, CSF2RB, PVALB, and PITX1, whereas the top genes downregulated in Sx included HBB, HBA2, SUCLA2, ANKRD2, and FHL1. Notably, several top-ranked genes were shared across the fibroadipogenic, pericyte/smooth muscle, and satellite/myogenic clusters, including ERBB3, PVALB, and MYH8 among genes upregulated in Sx, and HBA2 and HBB among genes downregulated in Sx. Because Visium spots capture mixed local transcriptomes, and because Ct and Sx biopsies were obtained from different quadriceps muscles, these shared signatures may partly reflect regional differences in local tissue composition and myofiber transcript contribution to non-myofiber spots. Additionally, because a substantial number of shared genes contributed to the pathway enrichment analyses, KEGG identified several overlapping pathways across clusters. The complete differential expression results for all clusters are provided in the supplemental **Table S2**.

### MuIS stratification identifies distinct transcriptional states in the Sx limb

To examine MuIS-associated differences in the periarticular muscle of the limb undergoing TKA, Sx samples were reintegrated and analyzed separately. Classification into MuIS-and MuIS+ groups was based on TNF-αR receptor expression (p = 0.0081; **Fig 3A**). UMAP analysis of the Sx-only dataset resolved spatial clusters corresponding to slow myofiber subpopulations (slow myofibers 1, 2 and 3), fast myofiber subpopulations (fast myofibers 1 and 2), fibroadipogenic cells, and pericyte/smooth muscle cells (**Fig 3B**). Canonical marker expression used for cluster annotations are shown in **Fig 3E**. When visualized by MuIS status, the major spatial domains were present in both groups without qualitative separation on the UMAP (**Fig. 3C**). Cluster composition analysis did not identify statistically significant differences between MuIS-and MuIS+ groups. Similar to our findings in the overall integration (Sx vs Ct) Sx-only integration shows that MuIS status is not associated with changes in the abundance of major spatial clusters.

**Figure 3.**
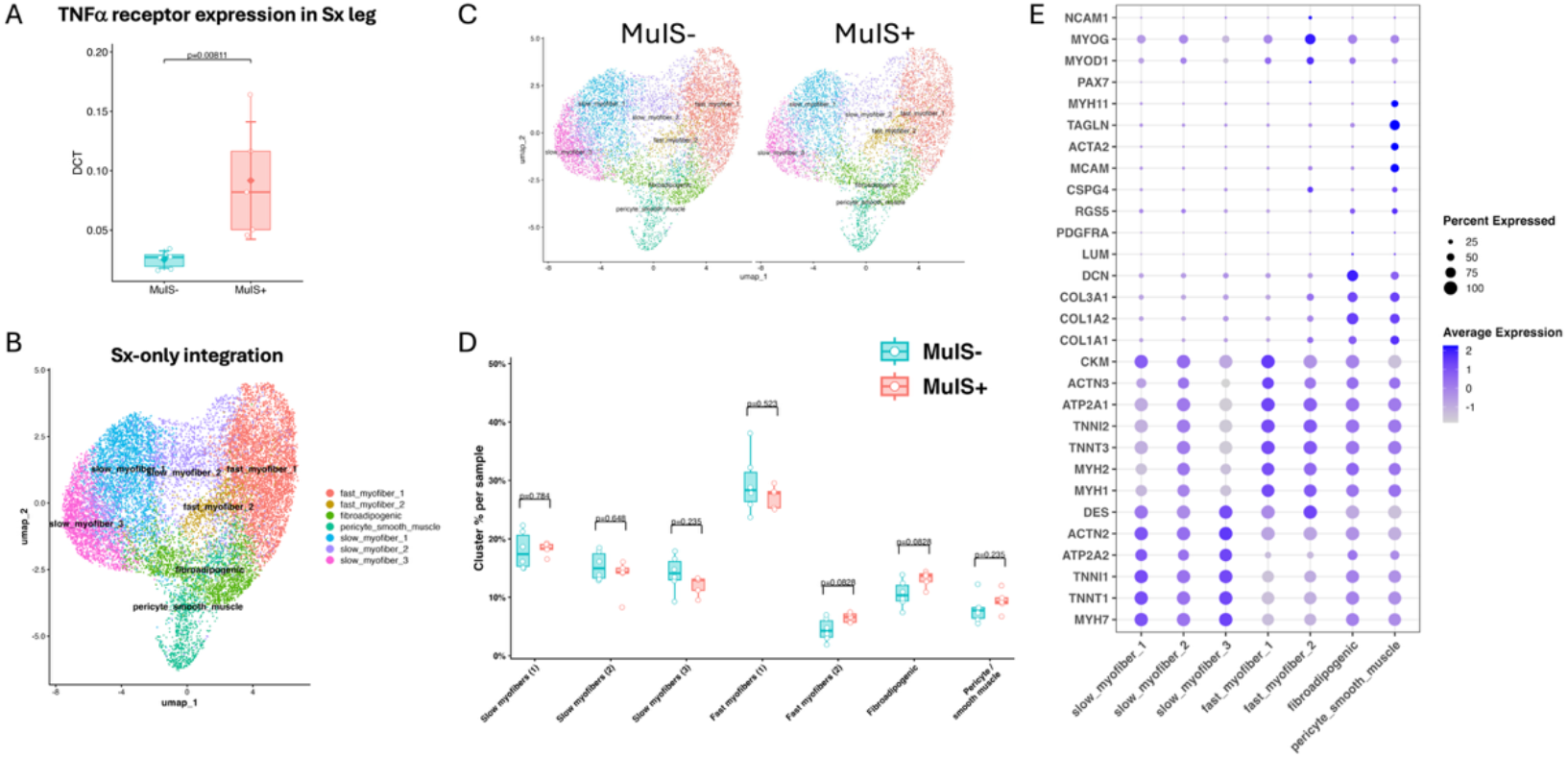
MuIS stratification in the surgical limb identifies distinct TNFα receptor expression states without major shifts in spatial cluster composition. (A) TNF-α receptor expression in the surgical (Sx) limb used to stratify participants into MuIS-and MuIS+ groups. Each point represents one participant and statistical significance was assessed using a Wilcoxon rank-sum test. (B) UMAP of the Sx-only integrated spatial transcriptomic dataset showing annotated clusters corresponding to slow myofiber subpopulations, fast myofiber subpopulations, fibroadipogenic cells, and pericyte/smooth muscle cells. (C) Split UMAPs of Sx samples stratified by MuIS-and MuIS+, showing the distribution of annotated clusters across MuIS groups. (D) Cluster composition per sample in Sx muscle, shown as the percentage of spots assigned to each annotated cluster in MuIS-and MuIS+ samples. Points represent individual samples, boxplots summarize sample distributions, and p values indicate group comparisons for each cluster. (E) Canonical marker gene dot plot used for annotation of Sx-only clusters. Dot size indicates the percentage of spots expressing each gene within a cluster, and color indicates average scaled expression.

Within-cluster DEG comparing MuIS+ vs. MuIS-identified broad transcriptional remodeling across all spatially defined clusters. The largest number of DEGs was observed in fast myofiber 1 (n = 490; 435 MuIS+ upregulated and 55 MuIS+ downregulated), followed by slow myofiber 1 (n = 420; 376 up and 44 down), slow myofiber 2 (n = 345; 285 up and 60 down), fibroadipogenic cells (n = 177; 151 up and 26 down), slow myofiber 3 (n = 161; 125 up and 36 down), fast myofiber 2 (n = 132; 109 up and 23 down) and pericyte/smooth muscle cells (n = 127; 99 up and 28 down). Across all clusters, the number of MuIS+ upregulated genes exceeded the number of MuIS+ downregulated genes. Interestingly. the slow-fiber compartment resolved into three subclusters with different slow-muscle signatures: slow myofiber 1 and 3 showed stronger MYH7 and TNNT1 expression, whereas slow myofiber 2 retained a weaker slow-marker profile with relatively more mixed fast-muscle signal. In contrast, the small satellite/myogenic cluster identified in the overall integration was not recovered as a distinct cluster in the Sx-only dataset, likely because of its low abundance and the smaller number of spots available after restricting the analysis to Sx-only.

In **Fig 4**, we focused on fast myofiber 1, fibroadipogenic, and pericyte/smooth muscle clusters because these populations are physiologically relevant to the KOA muscle pathology as discussed earlier. In fast myofiber 1, MuIS+ upregulated genes included MYH8, CHRNG, CHRNA1, LTK, and COL19A1, a pattern consistent with denervation-associated remodeling, re-expression of developmental myofiber, and local extracellular matrix adaptation. In contrast, MuIS+ downregulated genes included CRYM, the activity-responsive regulator NR4A3, and the transcripts IGHG1 and IGKC, in addition to the structural and contractile genes TPM2 and DES. This indicates loss of mature myofiber metabolic and contractile specialization and may be particularly relevant because downregulation of these has been linked to protein degradation and atrophy. In the fibroadipogenic cluster, MuIS+ similarly upregulated MYH8, COL19A1, LTK, CHRNG, and CHRNA1, suggesting that the fibroadipogenic spots are overlapping with denervation/remodeling spots. Downregulated genes included CRYM, IGHG1, NR4A3, MYL6B, and RSPO3, suggesting reduced stromal cell signaling, structural support, and maintenance of the local interstitial microenvironment. A similar pattern was observed in pericyte/smooth muscle cells, where MuIS+ upregulated MYH8, COL19A1, CHRNG, MYBPH, and MYL5, whereas CRYM, IGHG1, IGKC, RSPO3, and MYL6B were downregulated. Notably, as observed in the overall integrated analysis, several MuIS-associated DEGs were shared across clusters, suggesting a coordinated transcriptional remodeling across neighboring muscle compartments. Complete DEG results for all clusters are provided in **Table S3**.

**Figure 4.**
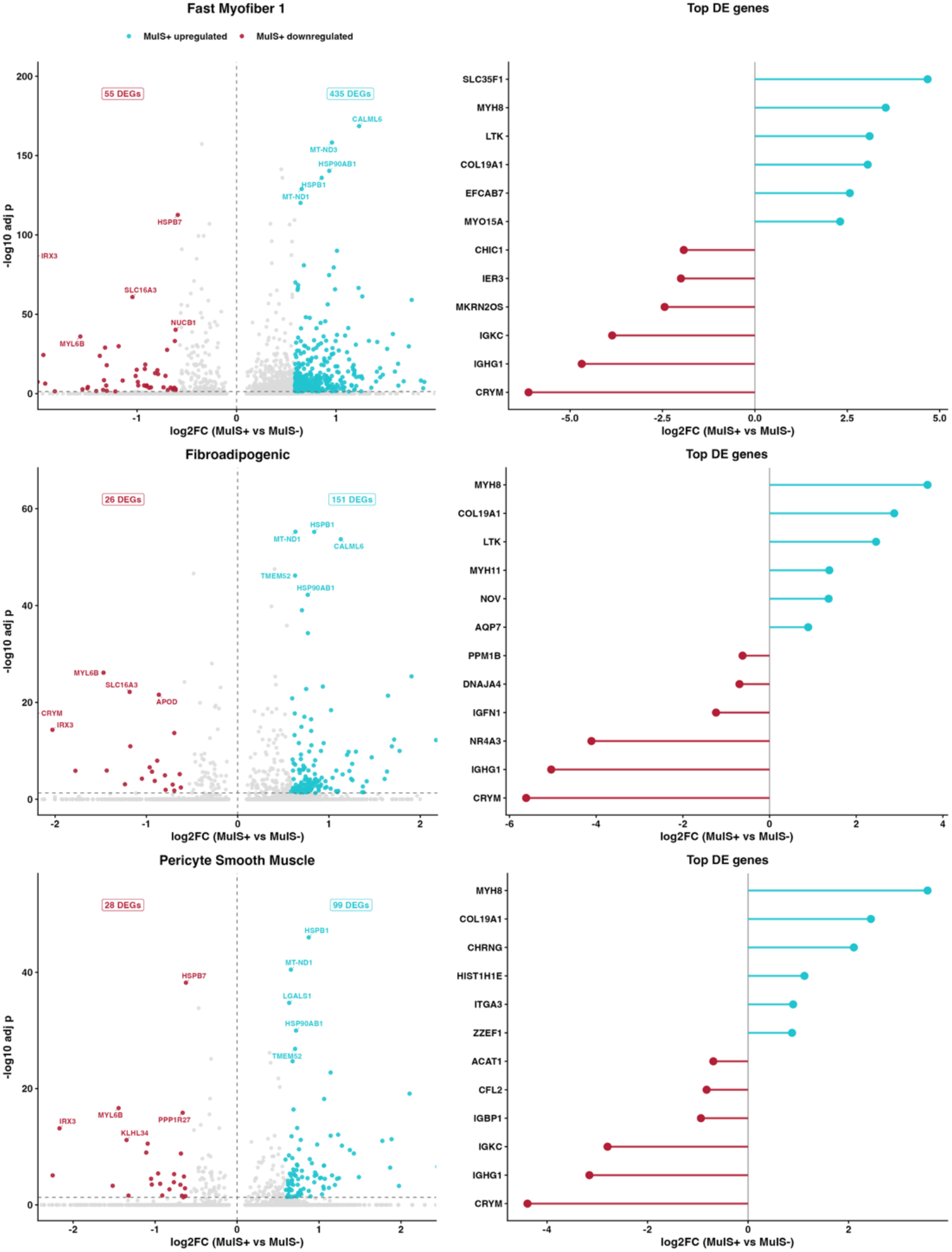
Within-cluster transcriptional differences between MuIS+ and MuIS-in surgical limb. Spatial transcriptomic spots from the Sx-only integration were stratified by MuIS status and differential expression was evaluated within annotated clusters. Shown are the three clusters prioritized for the main manuscript as they capture dominant myofiber transcriptional response (fast myofiber 1) and two biologically relevant interstitial populations involved in stromal remodeling and vascular support (fibroadipogenic and pericyte smooth muscle). For each cluster, the left panel shows a volcano plot of differential gene expression for MuIS+ vs. MuIS-spots, with selected top genes labeled. The right panel shows lollipop plots of upregulated and downregulated genes in MuIS+ within each cluster, ranked by differential expression significance and fold change. High-confidence differential expression was defined as adjusted p < 0.05 and absolute log2 fold-change of log2(1.5).

To determine whether these shared transcriptional features were accompanied by altered local tissue organization, we next performed neighborhood analysis of the Sx-only integration focusing on fast myofiber 1, fibroadipogenic, and pericyte/smooth muscle clusters (**Fig 5**). Because MYH8, CHRNG, CHRNA1, and MUSK were among the top MuIS+-enriched genes in fast myofiber 1 and are well recognized markers of denervation-associated and regenerative myofiber remodeling, we used their combined spatial expression to generate a denervation hotspot score. Representative spatial reconstructions showed that these three populations remained arranged in coherent local domains in both MuIS+ and MuIS-muscle, while denervation-associated hotspots were concentrated predominantly within fast myofiber 1 regions, with limited extension into adjacent stromal and vascular-support neighborhoods (**Fig 5A-B**). Pairwise neighborhood interaction analysis further showed that MuIS status did not markedly disrupt the overall spatial architecture of these compartments, as most cluster to cluster interaction z-scores were similar between groups (**Fig 5C**). Consistent with this, neighborhood enrichment heatmaps demonstrated strong self-enrichment within each cluster and relative depletion of cross-cluster adjacency in both MuIS-and MuIS+ samples (**Fig. 5D**).

**Figure 5.**
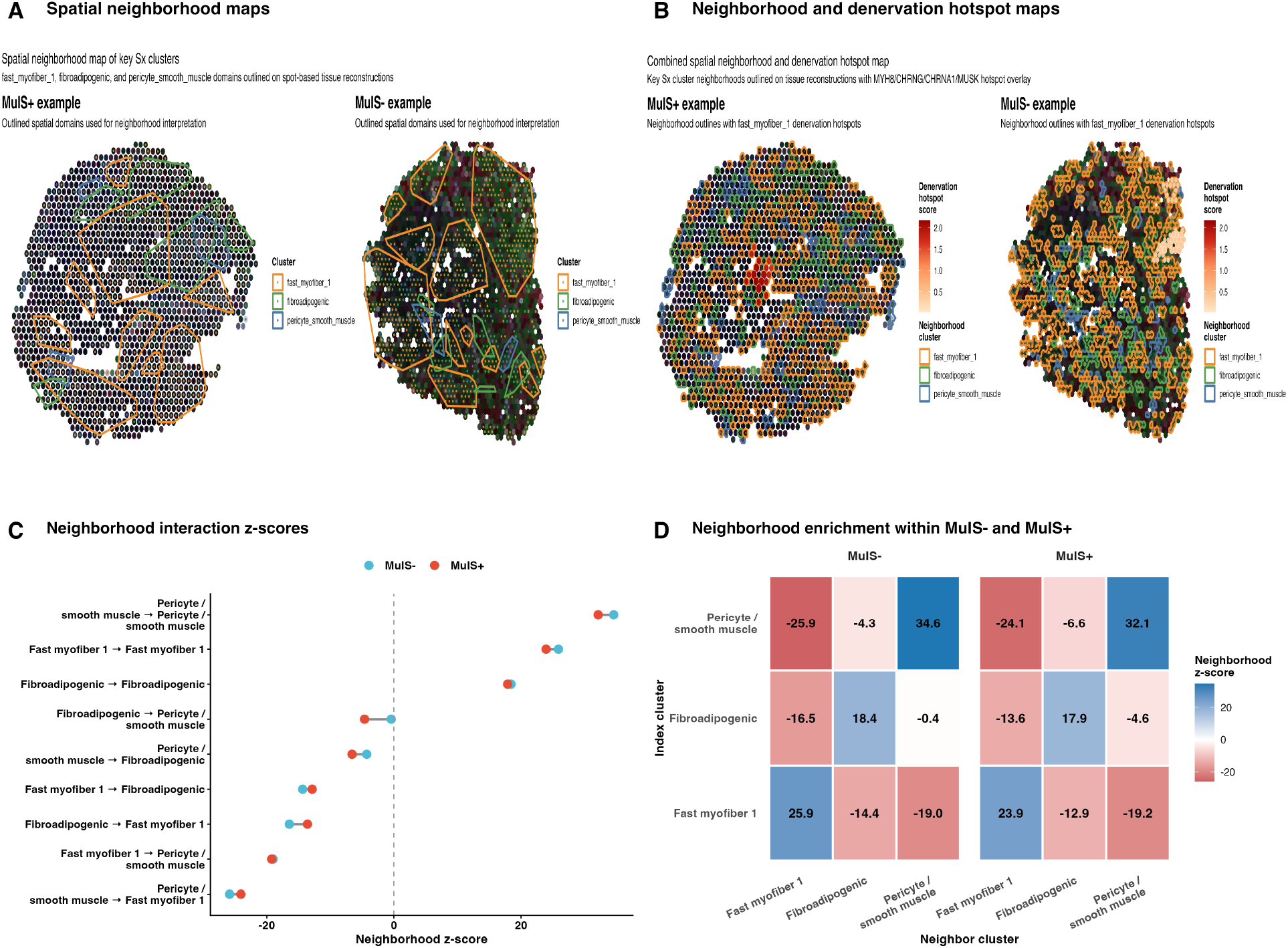
Spatial neighborhood organization and denervation-associated hotspot localization in MuIS-stratified surgical muscle. (A) Representative reconstructed spatial maps from MuIS+ and MuIS-surgical samples showing the distribution and local neighborhood relationships of the three key clusters fast myofiber 1, fibroadipogenic, and pericyte/smooth muscle. (B) Representative spatial reconstructions overlaid with denervation-associated hotspot scores derived from expression of MYH8, CHRNG, CHRNA1, and MUSK. (C) Pairwise neighborhood interaction z-scores comparing MuIS-and MuIS+ samples for the selected clusters. Positive values indicate enriched adjacency, whereas negative values indicate relative spatial separation. (D) Neighborhood enrichment heatmaps for MuIS-and MuIS+ samples s supporting broadly preserved local adjacency patterns, with strong self-enrichment within each cluster and relatively limited cross-cluster mixing.

**Figure 6.**
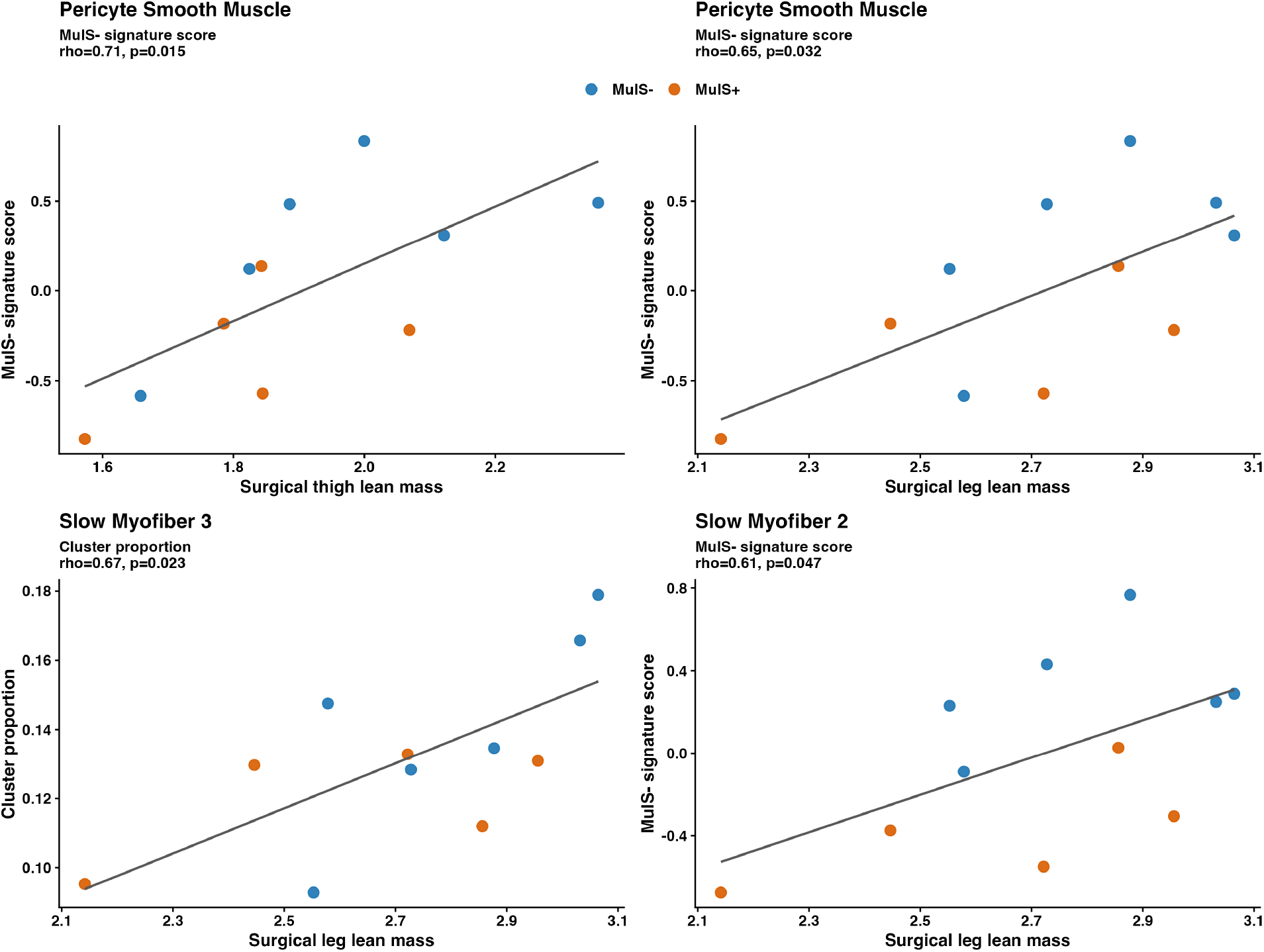
Top associations between Sx-leg spatial transcriptomic features and DXA-derived lean mass. Subject-level correlations were evaluated between features derived from the Sx-only spatial transcriptomic integration and normalized DXA measures of surgical leg or thigh lean mass. Each point represents one participant and is colored by MuIS status. Cluster proportion was defined as the fraction of Sx-leg spots assigned to a given cluster for that subject. MuIS-associated signature scores were calculated from subject-level pseudobulk data within each cluster as the mean z-scored expression of the top genes distinguishing MuIS+ and MuIS-. Associations were assessed using Spearman rank correlation, and the corresponding correlation coefficient (rho) and p value are shown in each panel.

Finally, to determine whether the MuIS-associated transcriptional program was specific to the surgical limb or also detectable in the control limb, we performed a Ct-only integration with de novo clustering followed by within-cluster DEG analysis comparing MuIS+ and MuIS-samples. The Ct-only dataset resolved clusters corresponding to multiple fast-and slow-myofiber subpopulations, two fibroadipogenic subclusters, and one pericyte/smooth muscle cluster (Fig. S1A). Applying the same differential expression criteria used in the main analyses, the largest Ct-only transcriptional effects were observed in slow myofiber 1 (n = 803; 739 MuIS+ upregulated and 64 MuIS+ downregulated), followed by fast myofiber 1 (n = 295; 217 up and 78 down), fast myofiber 2 (n = 248; 156 up and 92 down), fibroadipogenic 1 (n = 171; 113 up and 58 down), slow myofiber 2 (n = 160; 103 up and 57 down), fibroadipogenic 2 (n = 86; 26 up and 60 down), pericyte/smooth muscle (n = 47; 15 up and 32 down), and fast myofiber 3 (n = 10; 3 up and 7 down). Notably, the pericyte/smooth muscle cluster showed only modest transcriptional separation in Ct muscle, and the fibroadipogenic compartment split into two subclusters with distinct MuIS behavior. Full Ct-only within-cluster DEG results are provided in **Table S4**.

### Phenotype integration reveals lean mass-associated spatial programs in Sx limb

Exploratory integration of clinical phenotypes with the Sx-only spatial transcriptomic data suggested that muscle mass was more strongly related to cluster-level composition and transcriptional state than WOMAC-derived symptom measures. Specifically, higher surgical leg and thigh lean mass tended to associate with stronger MuIS-transcriptional programs in pericyte smooth muscle, and higher surgical leg lean mass also tended to associate with a greater proportion of slow myofiber 3 spots as well as stronger MuIS-associated expression in slow myofiber 2. These findings suggest that preservation of lean mass may align with retention of non-impaired transcriptional programs in vascular-support and slow-myofiber cell populations, whereas lower lean mass may be associated with a shift toward the MuIS+. As these analyses were performed in a small cohort these findings should be interpreted as exploratory and hypothesis-generating.

## DISCUSSION

We performed spatial transcriptomic profiling of skeletal muscle from individuals with end-stage KOA undergoing TKA to determine transcriptional states and how MuIS relates to local tissue remodeling. Several important findings emerged. First, the major annotated clusters were similarly represented in Sx and Ct limbs, indicating that advanced unilateral KOA was not associated with large shifts in the abundance of the dominant tissue compartments. Second, KOA-associated remodeling occurred primarily through transcriptional reprogramming within resolved clusters. Third, MuIS stratification identified a coordinated denervation and stress associated transcriptional program involving multiple clusters. These findings substantially extend our previous bulk RNA sequencing analyses of periarticular muscle, which identified MuIS as a clinically relevant molecular phenotype associated with inflammatory and catabolic signaling but could not resolve the origin of these transcriptional programs^7,10^.

### KOA Muscle Remodeling Occurs Within Preserved Spatial Domains

One of the major findings of the current study is that differences between Sx and Ct muscle were characterized primarily by transcriptional remodeling within spatially resolved clusters. This aligns with recent OA studies showing that relevant programs emerge across multiple joint tissues and cellular niches^3,22^. The overall clustering resolution was broadly preserved between Sx and Ct limbs despite advanced unilateral joint disease. We identified the expected major skeletal muscle domains, including slow and fast myofiber populations, fibroadipogenic cells, pericyte/smooth muscle cells, and satellite/myogenic cells. This was previously reported in human skeletal muscle single-cell and spatial atlases showing that adult skeletal muscle contains stable contractile, stromal, vascular, immune, and regenerative compartments^13,23^. However, we note that distinct endothelial and immune clusters were not recovered in the present dataset. This likely reflects the 55 µm spot resolution and the relatively low abundance of these populations, and it constrains our ability to localize inflammatory signaling.

Even at this lower resolution, the preservation of major spatial domains does not imply absence of disease-related remodeling. Instead, our paired-limb design suggests that KOA primarily alters the molecular state of resident cellular populations not their abundance. This distinction is important because skeletal muscle weakness, impaired muscle quality, and altered neuromuscular function are recognized contributors to KOA-related disability and incomplete recovery after TKA^4,24,25^.

Similarly, data from single-cell and spatial transcriptomics suggests that chronic diseases frequently arise through changes in existing cellular states. For example, single-cell analyses of osteoarthritic cartilage demonstrated that disease progression is accompanied by extensive transcriptional diversification of resident chondrocytes despite preservation of the major cellular populations^26^. Similarly, spatial transcriptomic profiling of osteoarthritic synovium identified localized inflammatory cell clusters that emerged through activation of resident fibroblast and immune transcriptional programs^22^. Additional data have also been reported in rheumatoid arthritis, where spatial transcriptomics demonstrated that pathological synovial architecture is largely maintained while disease progression is driven by coordinated transcriptional activation within existing stromal and immune niches^17^. Although not skeletal muscle, these studies support a molecular reprogramming of resident tissue compartments in KOA or similar conditions.

### Non-Myofiber Muscle Niches Remodeling in KOA

The high-confidence DEGs that remained after applying the 1.5-fold cutoff were distributed across myofiber and non-myofiber clusters, with particularly prominent signals in satellite/myogenic, fibroadipogenic, and pericyte/smooth niches (**Table S2**).These differences should also be interpreted considering the sampling design, as Ct biopsies were derived from vastus lateralis and Sx from vastus medialis (although no differences in fiber typing were found), therefore, part of the myofiber DEG may reflect the biology of the originating muscle in addition to disease-associated effects. Here we emphasize satellite/myogenic, fibroadipogenic and pericyte/smooth clusters because these non-myofiber niches are more physiologically informative for the KOA pathology, capturing stromal remodeling, extracellular matrix regulation, vascular support, and regenerative signaling within the periarticular muscle microenvironment.

Indeed, FAPs are essential regulators of extracellular matrix remodeling, muscle regeneration, and inflammatory signaling, while also contributing to the microenvironment required for efficient satellite cell activation and tissue repair^27^. Likewise, satellite cells, pericytes, endothelial cells, and other interstitial progenitor populations function as an integrated regenerative network whose coordinated activity determines skeletal muscle adaptation during aging, injury, and chronic disease^28,29^. Consequently, persistent transcriptional remodeling within these resident populations is expected to influence muscle quality long before overt alterations become apparent within mature myofibers.

On our pathway analysis, the contralateral limb retained pathways related to oxidative phosphorylation, thermogenesis, and other mitochondrial or contractile programs in the non-myofiber clusters highlighted in **Fig 2**. By contrast, the surgical limb showed enrichment of pathways associated with cytoskeletal remodeling and stress-responsive signaling, including focal adhesion, PI3K-Akt signaling, apoptosis, glucagon signaling, insulin resistance, and p53 signaling. The recurrence of similar pathways across clusters likely reflects overlap in the DEGs, together with the broad and redundant structure of KEGG pathway annotations, indicating shared local remodeling programs across these spatially adjacent non-myofiber clusters.

Our findings also reinforce observations from spatial transcriptomics on chronic muscle disorders, where pathological remodeling consistently localizes to specialized interstitial niches. Recent Visium-based analyses of muscular dystrophies demonstrated that fibroadipogenic progenitors accumulate within regions of active pathology and establish profibrotic cell-cell communication networks that precede extensive myofiber degeneration^16^. Similarly, single-cell spatial transcriptomics of facioscapulohumeral muscular dystrophy identified disease progression as a consequence of localized transcriptional reprogramming within regenerative and stromal cell populations instead of alterations across the entire muscle section^30^. Although OA is fundamentally distinct from primary myopathies, these studies support that chronic skeletal muscle pathology develops through remodeling of cellular neighborhoods instead of disruption of tissue architecture.

### MuIS+ reflects local transcriptional reprogramming in end-stage KOA

A major goal of this study was to determine whether the MuIS phenotype reflects differences in skeletal muscle organization or changes in the molecular state of resident populations. Following reintegration of the Sx-only dataset, MuIS+ and MuIS− samples retained remarkably similar features, with no significant differences in the abundance of major clusters. Our findings support the latter, demonstrating that, at the spatial resolution examined here, MuIS is primarily associated with transcriptional remodeling within resolved clusters.

The strongest MuIS-associated transcriptional effect in the Sx-only integration was observed in fast myofiber 1. In this cluster, MuIS+ led to a denervation/regeneration-associated program, with upregulation of MYH8 (11.6-fold), CHRNG (5.7-fold), CHRNA1 (5.1-fold), LTK (8.6-fold), and COL19A1 (8.3-fold). This pattern indicates re-expression of developmental myofiber (MYH8), altered neuromuscular junction signaling, and local extracellular matrix remodeling^31^. To our knowledge, this exact denervation/regeneration-like transcriptional signature has not been previously reported in human periarticular muscle from OA. However, our interpretation is supported by prior OA literature showing neuromuscular dysfunction and periarticular muscle remodeling^32^, as well as by preclinical KOA data demonstrating NMJ remodeling with altered acetylcholine receptor subunit and MuSK expression^33^. More broadly, the re-expression of MYH8, CHRNG, CHRNA1, and COL19A1 is an established denervation and regenerative pattern described in human unloading, aging, and denervation models^34–36^.

Using the combined spatial expression of MYH8, CHRNG, CHRNA1, and MUSK, we generated a composite denervation hotspot score and found that these hotspots were more prominent in MuIS+ muscle (**Fig 5B**). Interesting, denervation-associated signals were localized to discrete regions enriched within the fast myofiber 1 neighborhood. Prior spatial transcriptomic studies also showed that biologically coherent gene programs can localize to discrete tissue regions, including innervation-responsive domains in skeletal muscle and co-expression hotspots in other tissues^37–39^. This finding is further supported by our histological analysis, which showed higher median fiber shape index values in Sx, particularly in type 2 fibers, which represent a less regular and more angular fiber profile. Angular myofibers are a recognized feature of denervation-related remodeling and neurogenic atrophy, especially when present in scattered or grouped patterns within otherwise preserved tissue architecture^40^. However, fiber shape index did not differ between MuIS+ and MuIS− participants, potentially reflecting the limited statistical power of the Sx-only analysis given the small sample size (N = 11).

Transcriptional changes in fibroadipogenic and pericyte/smooth muscle clusters indicate that the MuIS+ is linked to stromal and vascular-support niches. As we discussed earlier, FAPs are a major source of extracellular matrix and can adopt persistent fibrogenic states in response to altered matrix stiffness, mechanical signaling, inflammation, and cellular senescence^41–43^. These findings also provide spatial context for our previous work defining MuIS in patients with end-stage KOA^10^. PLIER analysis previously identified endothelial, extracellular matrix, and fibroadipogenic latent variables (LVs) as major determinants of periarticular muscle inflammation but could not resolve their anatomical origin^7,10^. By preserving anatomical structure, the present spatial analysis shows that these disease-associated programs are embedded within discrete muscle microenvironments.

Since MuIS classification was originally based on whole-muscle TNF-αR expression, it is unlikely that the observed transcriptional phenotype originates exclusively from myofibers. Skeletal muscle contains a diverse mononuclear cell population, including FAPs^44^, endothelial cells and pericytes^45^, macrophages^46^, and other immune cells, all of which actively produce and respond to TNF-α signaling during tissue remodeling. In fact, macrophages are considered the predominant source of TNF-α in regenerating and chronically inflamed skeletal muscle, whereas FAPs, endothelial cells, and pericytes are major TNF-α responsive populations that amplify inflammatory and fibrogenic signaling through reciprocal interactions with immune and myogenic cells^29,47^. Accordingly, the absence of differential spot-level expression of TNF-α or its receptors in the spatial dataset does not contradict the whole muscle TNF-αR-based classification. At the 55-µm resolution used here, Visium spots capture mixed local transcriptomes and may have limited sensitivity for sparsely expressed transcripts. Instead, MuIS may reflect an integrated tissue-level TNF-responsive state whose downstream transcriptional consequences are distributed across multiple resident muscle compartments.

### Lean mass is associated with preservation of favorable spatial transcriptional programs

Although exploratory, our phenotype integration analyses suggest that lean mass may more closely reflect the biological state of periarticular skeletal muscle than patient-reported symptoms. Greater surgical-leg lean mass was associated with stronger MuIS− transcriptional programs within pericyte/smooth muscle and slow-myofiber compartments, whereas WOMAC scores demonstrated comparatively weak associations with spatial muscle features. These observations are biologically plausible once growing evidence indicates that muscle quality, rather than muscle quantity alone, is a major determinant of physical performance in KOA^4,48^. Additionally, a recent longitudinal study demonstrated that appendicular lean mass increases following TKA and parallels improvements in musculoskeletal health^49^.

This study is not without limitations. First, Ct samples derived from vastus lateralis and Sx samples from vastus medialis. Therefore, some limb-related transcriptional differences may reflect intrinsic biology in addition to disease-associated remodeling. This concern is mitigated in the Ct-only and Sx-only integrations used for MuIS stratified analyses and our fiber typing analysis showing no differences between the two muscles. Second, MuIS stratification resulted in an imbalanced cohort composition, particularly with limited representation of women in the MuIS+ group, which constrains interpretation of potential sex-specific effects. Third, spatial transcriptomic spots capture mixed local transcriptomes, and therefore cluster assignments reflect dominant regional signatures rather than fully resolved individual cell states. Despite these limitations, the paired-limb design and spatially resolved approach provide a useful framework for identifying biologically meaningful muscle phenotypes in end-stage OA. To our knowledge, this is the first spatial transcriptomic analysis of periarticular skeletal muscle in patients with end-stage KOA and likely one of the largest spatial transcriptomic skeletal muscle datasets reported to date.

In conclusion, we show that periarticular skeletal muscle in end-stage KOA retains similar representation of major annotated cellular compartments, while the dominant differences arise through localized transcriptional remodeling within specific muscle-resident populations. The finding that MuIS is associated with coordinated transcriptional remodeling across myofiber and interstitial domains, including a prominent denervation-and regeneration-associated program localized primarily to fast myofiber regions, supports that MuIS reflects a distinct molecular state within periarticular skeletal muscle. These findings provide a physiologically relevant framework for understanding muscle dysfunction in KOA and identify spatially resolved transcriptional programs as potential targets for patient stratification and future interventions aimed at improving recovery after TKA.

## Acknowledgments and funding.

Parent grant R01HD084124. Analyses for this follow-up paper were supported by internal funds at Regeneron Pharmaceuticals and the Florida Institute for Human and Machine Cognition.

---

**Supplemental Figure 1.**
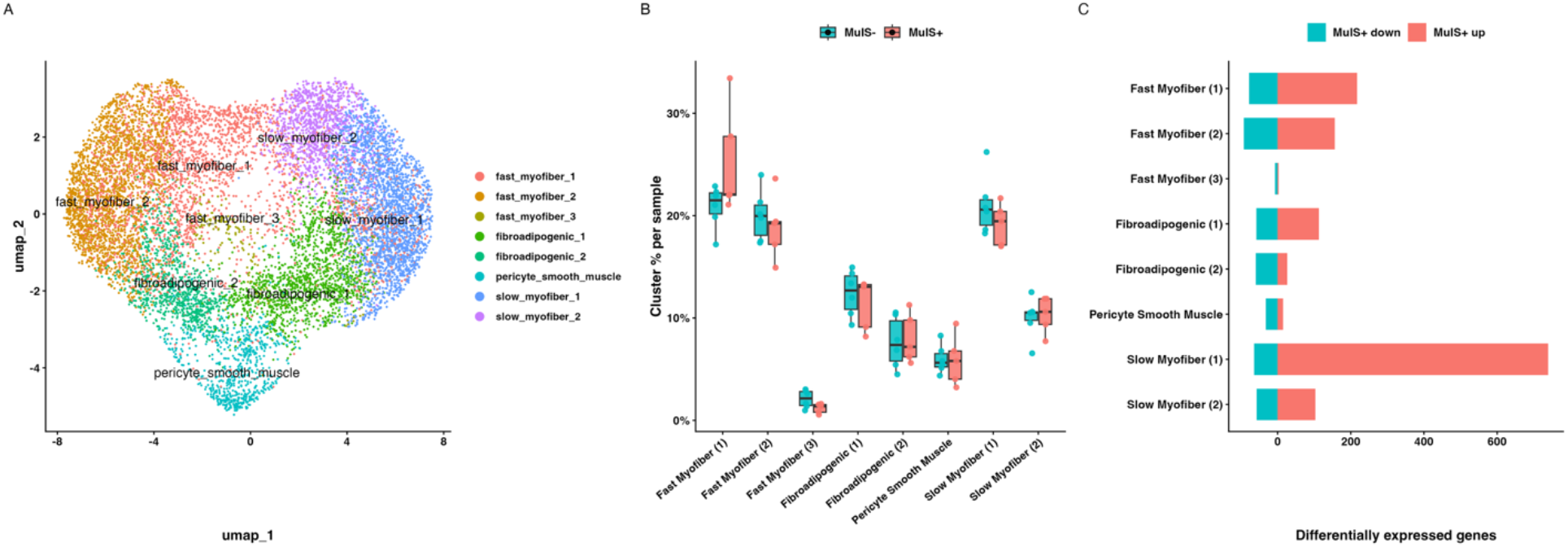
Ct-only integration identifies a weaker and less coherent MuIS-associated transcriptional program than the surgical limb. (A) Annotated UMAP of the Ct-only integration showing de novo clustering of control-leg spatial transcriptomic spots into fast-and slow-myofiber subpopulations, two fibroadipogenic subclusters, and one pericyte/smooth muscle cluster. (B) Sample-level cluster composition by MuIS status in the Ct leg, showing broadly similar representation of major cell populations between MuIS+ and MuIS-samples. (C) Summary of within-cluster differential expression in the Ct-only analysis comparing MuIS+ vs. MuIS-, displayed as the number of MuIS+ upregulated and MuIS+ downregulated genes per cluster.

